# Redox-activated induction of Warburg-type metabolism in the adult heart

**DOI:** 10.64898/2026.02.18.706512

**Authors:** Youwen Yang, Adam A Nabeebaccus, Olga Mikhaylichenko, Christina M Reumiller, Domenico Cozzetto, Asjad Visnagri, Anna Zoccarato, Min Zhang, Alison C Brewer, Anne D Hafstad, Celio CX Santos, Ajay M Shah

**Affiliations:** King’s College London British Heart Foundation Centre, School of Cardiovascular and Metabolic Medicine & Sciences, London, UK

## Abstract

Proliferating cells rewire glucose metabolism away from oxidative ATP production towards glycolysis and anabolic branch pathways (the Warburg effect). In contrast, the contractile function of terminally differentiated cardiomyocytes in the heart critically depends upon mitochondrial ATP generation. Adult cardiomyocytes are largely non-proliferative but undergo hypertrophy in response to increased heart workload. A fundamental question in the field is how cardiac metabolism is modified to balance competing energetic and anabolic demands. We previously reported that the redox-signalling H_2_O_2_-generating enzyme, NADPH oxidase 4 (NOX4), facilitates compensated cardiac function in hearts undergoing hypertrophic remodelling. Here, we show that NOX4 induces Warburg-type reprogramming of glucose metabolism in the healthy heart, with increased flux into the pentose phosphate, serine and nucleotide biosynthetic pathways. Using an integrated multiomics approach, we uncover a NOX4-regulated network involving interplay between direct transcriptional activation of metabolic genes via NRF2 [aka NFE2L2] and ATF4 and a broader epigenetic regulation. This reprogramming of glucose intermediary metabolism along with previously reported NOX4-mediated enhancement of fatty acid oxidation may serve to optimally support the dual requirements of increased energy demand and remodelling in the heart. Our findings identify a novel paradigm for redox-regulated Warburg-type metabolic reprogramming in the terminally differentiated heart.

## Main

The continuous contractile activity of the adult heart is driven by terminally differentiated cardiomyocytes with minimal proliferative capacity and depends critically upon ATP generated from the oxidation of carbon-based nutrients such as fatty acids, glucose and ketone bodies.^1,2^ Physiological stresses such as pregnancy and chronic exercise and pathological stresses such as an increased haemodynamic load induce heart enlargement via hypertrophic remodelling, supporting increased cardiac work. Whereas contractile function is maintained during physiological remodelling, persistent pathological remodelling usually leads to contractile dysfunction and heart failure.^3^ The contribution of alterations in energetic pathways (e.g. utilisation of different energy sources) to heart dysfunction has been a predominant focus of research to elucidate underlying pathophysiology. Less attention, however, has been paid to the metabolic pathways required to support heart remodelling, such as the biosynthesis of cellular building blocks.^4^ It is suggested that the channelling of carbons derived from glucose into ancillary biosynthetic pathways is important in the developing heart and during chronic exercise but few studies have directly assessed metabolic flux. Most importantly, it remains unclear how the metabolic changes to support an increase in biomass in the adult heart are coupled to alterations in energy production and how the overall response is regulated.

NOX4 is an H_2_O_2_-generating enzyme that is induced in the heart in response to chronic haemodynamic overload or other stresses and promotes adaptive hypertrophic remodelling with better preserved contractile function.^5-7^ The actions of NOX4 in the heart and other tissues involve specific redox signalling, such as the activation of the transcription factors NRF2 and ATF4 or an augmentation of Akt signalling.^7-11^ Prompted by an analysis of the NOX4-modulated cardiac proteome, we previously identified that NOX4 induces a perturbation of myocardial glucose metabolism with a shift away from glucose oxidation to fatty acid oxidation and an overall preservation of bioenergetics as assessed by ^32^P-NMR spectroscopy.^12^

To define the wider metabolic alterations induced by NOX4, we profiled the total abundance of >150 metabolites in pathways linked to glucose intermediary metabolism in the hearts of mice with cardiac-specific overexpression of NOX4 (csNOX4TG) and matched wild-type littermate (WT) hearts. There were marked differences between the two groups with notably higher pools of cystathionine, glutathione, glycolytic intermediates, phosphoserine, nucleotide precursors and many other metabolites in csNOX4TG hearts (Fig. 1a, Extended data Fig.1a, Supplementary Table 1). A metabolite set analysis revealed perturbation of several ancillary glycolytic and anabolic pathways, the TCA cycle, glutathione metabolism, and a signature for the Warburg effect^13^ (Fig. 1b). To study glucose-dependent intermediary metabolic pathways in more detail, we undertook mass spectrometry-based stable isotope-resolved metabolomics. Isolated csNOX4TG and WT hearts were perfused with uniformly labelled glucose (U^13^C-glucose) and ^13^C-label incorporation into downstream metabolites was quantified (Fig. 1c-j). This revealed a significantly higher % incorporation of ^13^C-label into intermediates in the pentose phosphate pathway (PPP; Fig. 1e), hexosamine biosynthesis pathway (HBP; Fig. 1f), serine biosynthesis pathway (SBP; Fig. 1g) and nucleotide biosynthesis (Fig. 1h) in csNOX4TG hearts. We observed a lower % ^13^C incorporation into glycolytic metabolites (Fig. 1d), TCA cycle intermediates (Fig. 1i) and aspartate (Fig. 1j) in csNOX4TG. The comparative abundance of ^13^C-labelled and ^12^C metabolites is shown in Extended data Fig. 1b-h. These results suggest an increased flux of glucose-derived carbons into several glycolytic branch pathways in csNOX4TG hearts along with a corresponding reduction in flux into the TCA cycle. While we did not detect intermediates of glycogen biosynthesis, absolute levels of glycogen were significantly higher in csNOX4TG hearts compared to WT (Extended data Fig.1i). We also observed higher levels of intermediates in the serine-glycine-one-carbon (SGOC) pathway in csNOX4TG (Supplementary Table 1).

**Fig. 1.**
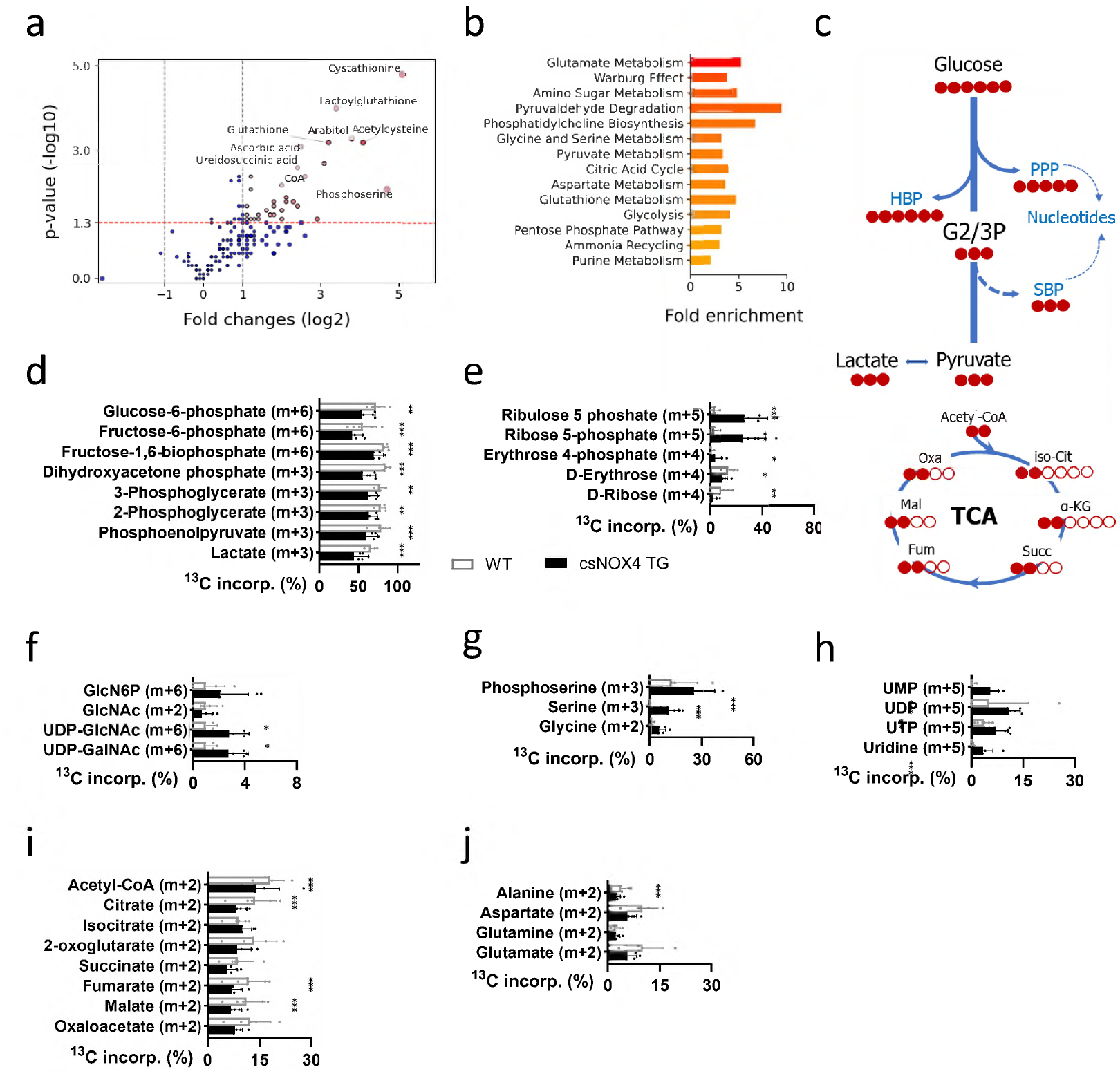
NOX4 induces widespread rewiring of glucose intermediary metabolism in the heart. **a**, Volcano plot depicting changes in relative metabolite abundance in csNOX4TG (n=4) vs. WT (n=3) hearts analysed using MetaboAnalyst 6.0. Horizontal dotted red line indicates the p value significance threshold while vertical dotted lines show the fold change (FC) thresholds. Key metabolites including cystathionine, glutathione, glycolytic intermediates, phosphoserine, and nucleotide precursors are labelled. **b**, Metabolite set enrichment (MetaboAnalyst) showing main metabolic pathways enriched in csNOX4TG vs WT. **c**, Schematic of ^13^C labelling of metabolites after heart perfusion with [U-^13^C_6_]glucose. PPP, pentose phosphate pathway; HBP, hexosamine biosynthetic pathway; G2/3P, glucose 2,3 biphosphate; SBP, serine biosynthetic pathway; TCA, tricarboxylic acid cycle; iso-Cit, isocitrate; α-KG, α-ketoglutarate; Succ, succinate; Fum, fumarate; Mal, malate; Oxa, oxaloacetate. Filled circles indicate ^13^C, empty circles indicate ^12^C. **d-j**, Percentage ^13^C incorporation in metabolites from hearts perfused with [U-^13^C_6_]glucose. Data are mean ± SEM for wild-type (WT, n=5) and csNOX4Tg (n=7) groups. Metabolites shown are from glycolysis (d), PPP (e), HBP (f), SBP (g), nucleotide metabolism (h), TCA cycle (i) and amino acids (AA) (j). GlcN6P, glucosamine 6-phosphate; GlcNAc, N-Acetylglucosamine, GalNAc, N-acetylgalactosamine. Filled bars denote the ^13^C isotopologue with the specific moiety shown in brackets. Comparisons were performed using Mixed Model analysis; *p≤0.05, **p≤0.01, ***p≤0.001.

Taken together, these data indicate that NOX4 induces a widespread remodelling of glucose intermediary metabolism in the adult heart, with a channelling of glucose carbons into anabolic and other glycolytic branch pathways and away from the TCA cycle, analogous to Warburg-type reprogramming in proliferating tissues.^13^

To investigate potential mechanisms underlying these NOX4-induced metabolic changes, we undertook RNA-seq analyses of csNOX4TG and WT hearts. This showed 2230 upregulated genes and 742 downregulated genes in csNOX4TG compared to WT hearts (Fig. 2a). We performed functional enrichment analyses to obtain a global view of the pathways linked to differentially expressed genes (DEGs), focusing on metabolic and signalling pathways. The top metabolic pathways enriched in csNOX4TG included pyrimidine ribonucleotide synthesis, PPP, serine, glycine and glutathione metabolism (Fig. 2b), suggesting that a significant component of the changes observed by metabolomics may be transcriptionally mediated. There was also significant enrichment of other pathways linked to anabolism including tRNA charging (protein synthesis), superpathway of inositol phosphate compounds (involved in glycerophospholipid synthesis) and dolichyl-diphosphooligosaccharide (*N*-glycan) biosynthesis. The most downregulated metabolic pathways by z-score analysis included ketone body and branch chain amino acid metabolism (Extended data Fig. 2c). The top enriched signalling pathways by z-score in csNOX4TG hearts included NRF2-mediated stress response and EIF2 signalling (Fig. 2c), pathways previously shown to be specifically activated by NOX4.^7-10^ A more focused gene set enrichment analysis of the key metabolic pathways identified in the above analysis along with the validation of changes in expression of selected genes by RT-PCR is shown in Fig. 2d-k and Extended data Fig. 2d.

**Fig. 2.**
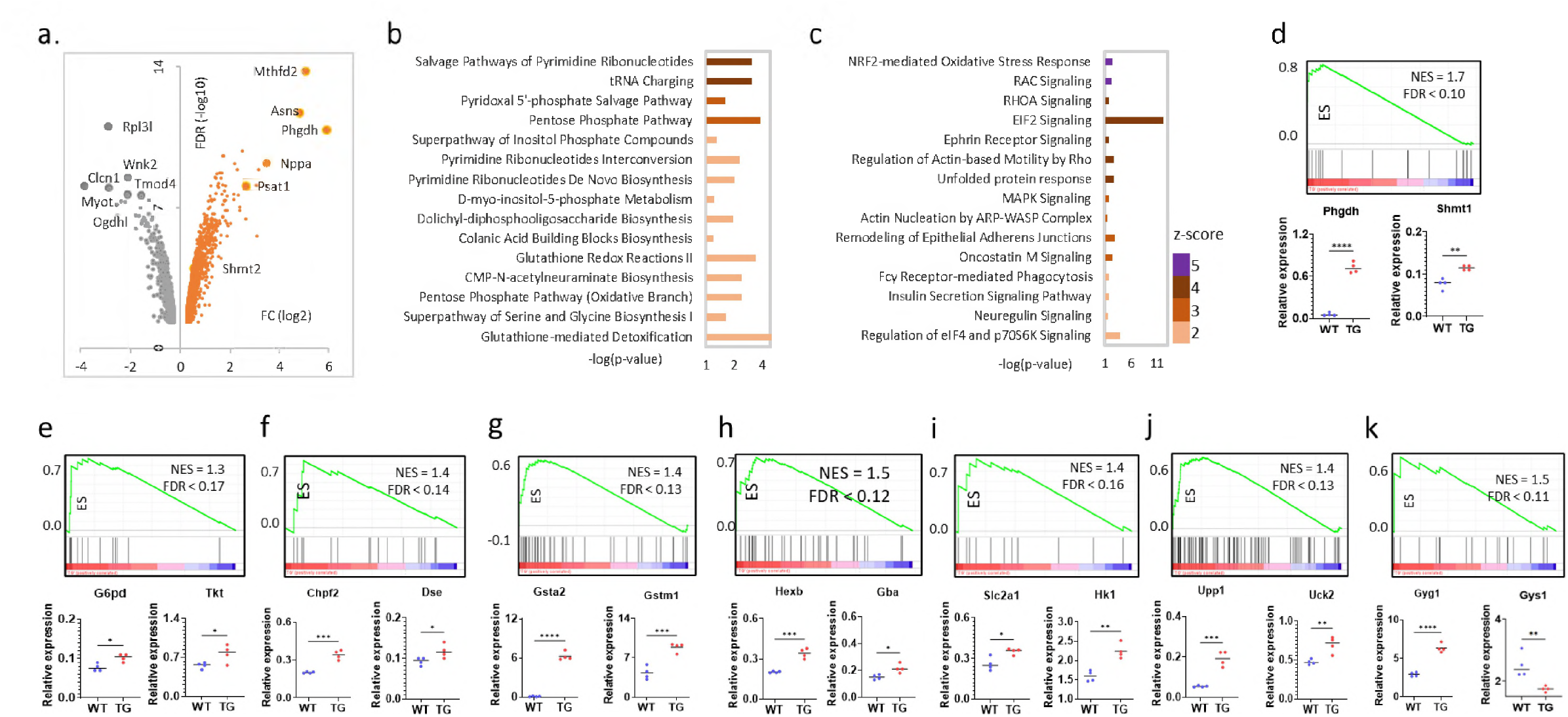
Effects of NOX4 on the cardiac transcriptome. **a**, Volcano plot depicting differentially expressed genes in RNA-seq data from csNOX4TG and WT hearts (FDR<0.05, gene expression fold change [FC] >1.2 or <1.2). *N*=4 biological replicates per group. **b**-**c**, The top metabolic and signalling pathways (b and c, respectively) enriched in csNOX4TG hearts, as analysed using IPA software (Qiagen); a z-score of ≥ 2 is used to indicate significantly activated pathways. **d-k**, Gene set enrichment analysis (GSEA) of metabolic pathways derived from the RNA-seq data, by enrichment score (ES) ranking. Validation by RT-qPCR of key selected genes from the GSEA analyses is shown below the GSEA plots: **d**, glycine, serine & threonine metabolism. **e**, pentose phosphate pathway. **F**, glycosaminoglycan biosynthesis. **g**, glutathione metabolism. **h**, glycosphingolipid metabolism. **i**, aerobic glycolysis. **j**, nucleotide metabolism. **k**, glycogen metabolism. n = 4 independent experiments. p-values were calculated using an unpaired t-test; *p ≤ 0.05, **p ≤ 0.01,***p ≤ 0.001.

Potential upstream transcriptional regulators of the changes in gene expression were investigated using IPA software (Qiagen). This suggested that NRF2, ATF4, c-MYC, MLXIPL (ChREBP) and p53 were amongst several transcription factors predicted to be activated based on the overlap between z-score ranking and p-value (Extended data Fig. 2a). Conversely, KLF15 and PPARGC1a were among the top transcriptional regulators predicted to be repressed (Extended data Fig. 2b). Other transcriptional regulators predicted to be repressed included SMARCA5, a protein that forms part of a chromatin remodelling complex, and GPS2, a core part of the NCOR1-HDAC3 corepressor complex. These results indicate a broad transcriptional regulation of intermediary metabolic pathways by NOX4, characterised by a complex interplay among several transcription factors known to modulate cellular metabolism and proteins involved in chromatin accessibility.

Among these transcription factors, ATF4 and NRF2 are known to be directly and specifically activated by NOX4.^7-10^ We performed chromatin immunoprecipitation followed by quantitative PCR (ChIP-qPCR) in csNOX4TG and WT hearts to validate the direct transcriptional regulation of selected key genes by these factors (Fig. 3a). We confirmed enhanced ATF4 binding at the promoter regions of genes encoding key enzymes in the SBP (*Phgdh*), one-carbon metabolism (*Shmt2*), amino acid transport (*Slc1a4*), and nucleotide metabolism (*Txnrd1*) in csNOX4TG hearts, while NRF2 showed increased binding to promoters of antioxidant defence genes (*Gclm, Gsr*) and the PPP (*G6pd*). These data confirm that NOX4-mediated metabolic reprogramming at least in part involves the direct binding of NRF2 and ATF4 to metabolically relevant gene loci.

**Fig. 3.**
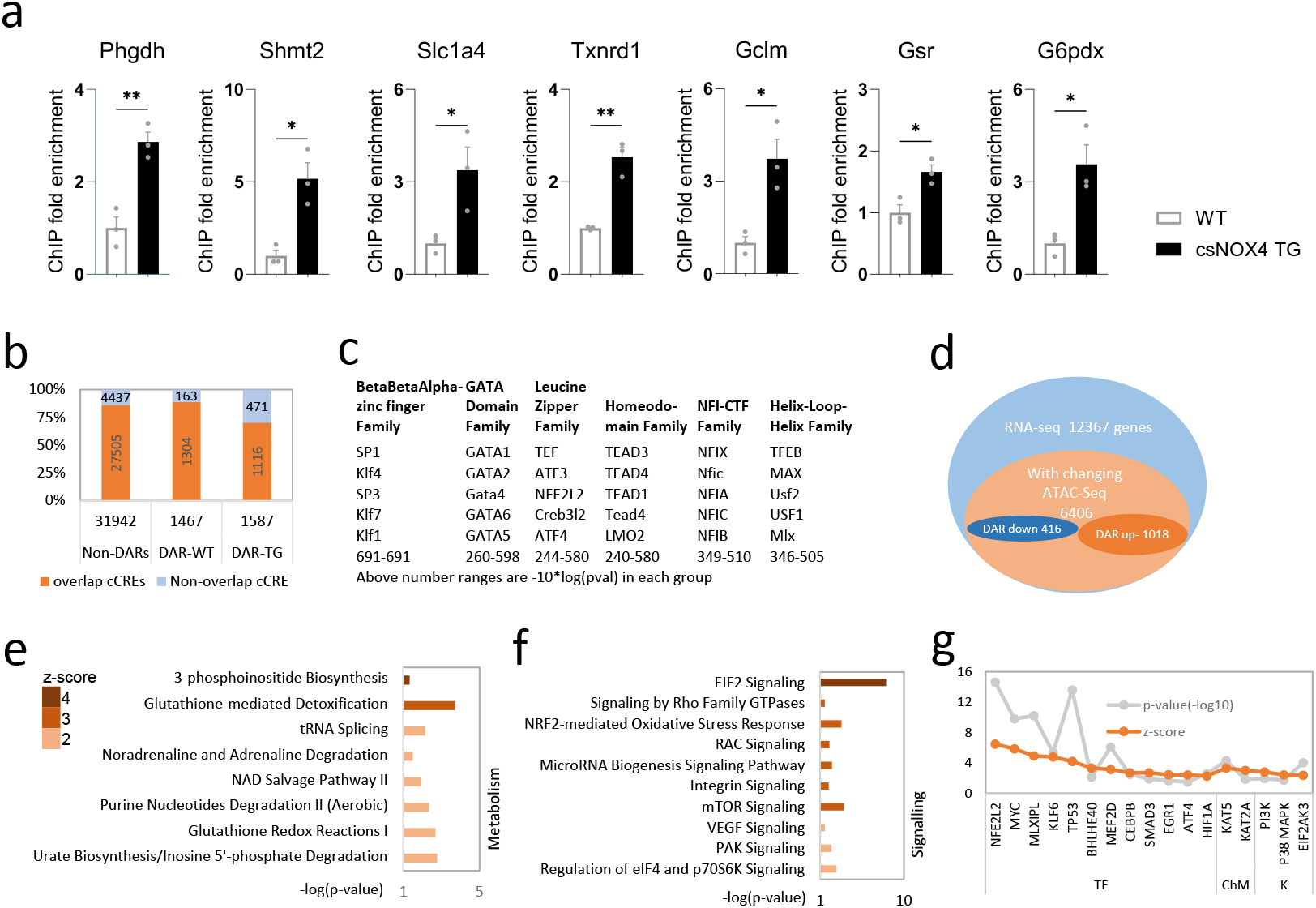
NOX4 increases chromatin accessibility for genes regulating metabolic and stress-responsive pathways. **a**, ChIP-qPCR analyses showing the enrichment of ATF4 at gene loci for *Phgdh, Shmt2, Slc1a4* and *Txnrd1*, and of NRF2 at gene loci for *Gclm, Gsr* and *G6pdx*. Data are presented as fold enrichment relative to WT samples, after normalisation to both the respective input control and a gene desert region. Mean ± SEM of three independent experiments; *P < 0.05, **P < 0.01 by Student’s t-test. **b**, ATAC-seq peaks were classified into three group: at open chromatin within Differential Accessibility Regions (DARs) in TG or WT, and non-DARs. The proportion of peaks that overlapped with known candidate cis-regulatory elements (cCREs) are shown for each group. ATAC-seq peaks were called using MACS2 (v. 2.2.6) with the default q-value threshold of 0.05. The DARs between WT and TG groups were evaluated using the Wald test as implemented in DESeq2 (n=4 samples, each group). cCRE data was downloaded from UCSC Genome Browser. **c**, Top enriched transcription factor family DNA binding motifs identified within the DARs of open chromatin in TG hearts, ranked by lowest p-values. Values shown at the bottom are -10*log(p-value); e.g., p = 0.01 and p = 0.001 correspond to values of 20 and 30, respectively. Analysis used the Seqpos motif tool in the Cistrome toolbox, with default parameters (http://cistrome.org/ap/). **d**, Integration of RNA-seq and ATAC-seq datasets. Venn diagram shows the overlap between RNA-seq detected genes (12,367 genes, blue circle) and genes associated with DARs from the ATAC-seq analysis (6,406 genes, orange circle). Within the latter group, 1,018 genes were upregulated in expression and showed increased accessibility (DAR up) in TG hearts while 416 genes were downregulated in expression and showed decreased accessibility (DAR down). **e-f**, IPA analysis of the 1,018 upregulated genes with increased chromatin accessibility (DAR up) for enrichment of metabolic (e) or signalling (f) pathways. **g**, Analysis of predicted upstream transcriptional regulators for the upregulated genes associated with open chromatin at DARs in TG hearts. TF, transcription factor; ChM, chromatin modifier; K, kinase. Enrichment cut offs used were a z-score ≥ 2 and a -log(p-value) >1.3.

We next undertook ATAC-sequencing to investigate how NOX4 influences chromatin accessibility and to gain a genome-wide perspective on regulatory networks underlying the RNA-seq results. Around 35,000 unique accessible regions (peaks) were detected across csNOX4TG and WT hearts of which approximately 3,000 were differentially accessible regions (DARs) (Fig. 3b). 70% of open DARs in csNOX4TG hearts overlapped with known candidate *cis*-regulatory elements (cCREs) as compared to 89% of open DARs in WT hearts (Fig. 3b), suggesting that NOX4 increases the proportion of uniquely regulated open chromatin regions. Analysis of the top transcription factor family motifs unique to the NOX4 group revealed association with a large number of factors including SP1, NRF2, ATF4 and members of the KLF, GATA, NF1 and TEAD families (Fig. 3c).

To understand the effects of changes in chromatin accessibility at a transcriptional level, we performed an integrated analysis of the RNA-seq derived DEGs and the ATAC-seq DARs (Fig. 3d). Pathway analysis of the up-regulated gene set showed a strong enrichment of key metabolic and signalling pathways including glutathione metabolism, tRNA splicing, EIF2 signalling, NRF2 response and mTOR signalling (Fig. 3e-f). Furthermore, an analysis of the predicted upstream transcriptional regulators showed that NRF2, followed by c-MYC, KLF6 and MLXIPL were the most significantly activated by z-score (Fig. 3g), supporting the idea that open chromatin regions of genes regulated by these transcription factors in part drive the upregulated gene expression. Non-cCRE-associated genes within DARs identified in this analysis are listed in Extended data Fig. 3c. We also identified a small number of downregulated metabolic pathways (associated with closed DARs) that were enriched in csNOX4TG hearts (Extended data Fig. 3a). Putative transcription factors and regulators that may contribute to these pathways are shown in Extended data Fig. 3b and included KDM5a, a histone demethylase known to be a key regulator of the epigenome (including metabolic genes).^14^ Taken together, these results suggest that genome-wide transcriptional effects of NOX4 are orchestrated largely at the chromatin level.

Previous work from our group and others uncovered that NOX4 specifically promotes activation of the transcription factors NRF2 and ATF4.^7-10^ The results presented so far suggest that while these transcription factors contribute significantly to the metabolic alterations in csNOX4Tg hearts, they do not account for all the changes as numerous genes regulated by many other transcription factors are also perturbed. Since epigenetic mechanisms are well established as being metabolically regulated,^15^ we investigated whether they also contribute to the NOX4-mediated genetic programme. We focused on histone and DNA methylation given that our metabolomics analyses suggested altered activity of the SGOC pathway (Supplementary Table 1) which generates S-adenosyl methionine required for DNA and histone methylation;^16^ and reduced carbon flux into TCA cycle intermediates where α-ketoglutarate is a co-factor in demethylation reactions.^15^ A genome-wide CUT&Tag analysis of histone H3K4me3 marks revealed that while csNOX4TG hearts showed fewer differentially enriched regions than WT hearts, they exhibited a substantially higher number of unique H3K4me3 peaks at both promoters and distal promoter regions (Fig. 4a). Analysis of differentially expressed genes associated with differential or unique H3K4me3 peaks showed predominant upregulation in csNOX4TG hearts (Fig. 4b), with significant enrichment in metabolic and signalling pathways (Fig. 4c-d). While the enriched pathways aligned with those identified in our RNA-seq data, H3K4me3 analysis revealed a broader set of predicted upstream transcriptional regulators (Fig. 4e), suggesting that H3K4me3 deposition may precede or mark poised genes that drive the transcriptional program. Notably, H3K4me3 enrichment was observed at promoters of key metabolic genes spanning pathways involved in the PPP, one-carbon metabolism, redox homeostasis and lipid metabolism.

**Fig. 4.**
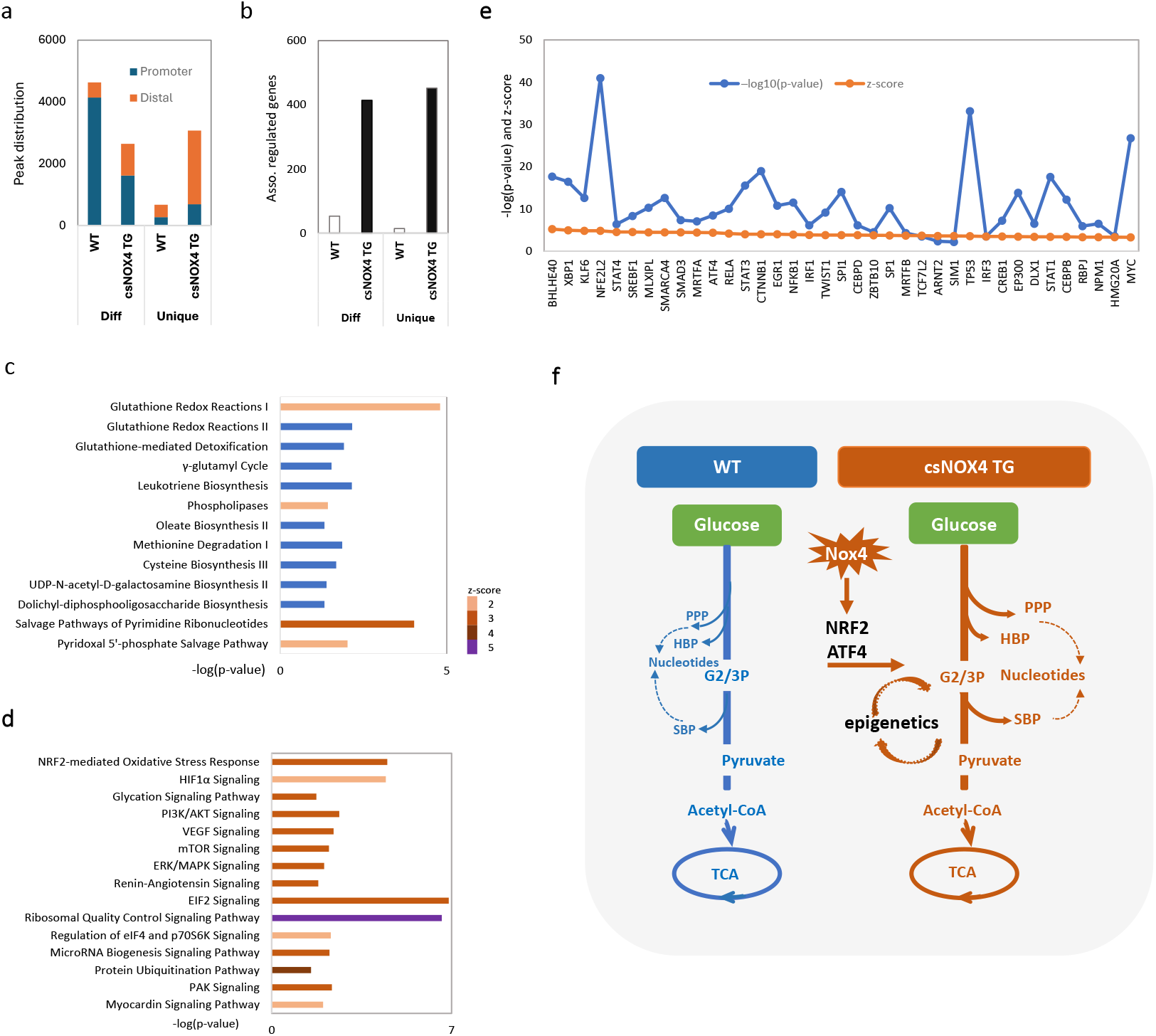
Effects of NOX4 on cardiac H3K4me3 and DNA methylation. **a**, Number of enriched differential binding regions (Diff) and unique H3K4me3 peaks in WT and csNOX4TG hearts at promoters and distal regions. **b**, Number of upregulated genes associated with enriched Diff or unique H3K4me3 peaks in WT and csNOX4TG. **c-d**, Top metabolic pathways (c) and signalling pathways (d) enriched in csNOX4TG hearts, as analysed using IPA software; a z-score of ≥ 2 is used to indicate significantly activated pathways. Blue columns denote pathways in which the IPA software could not determine a z-score. **e**, Top upstream transcription regulators predicted to be activated. Enrichment cut offs used were a z-score ≥ 2 and a -log(p-value) >1.3. **f**, Proposed model for NOX4-driven metabolic reprogramming. NOX4 specifically activates NRF2 and ATF4 and thereby increases glucose carbon flux through ancillary glycolytic pathways. Wider metabolic reprogramming by NOX4 involves epigenetic regulation.

We also assessed whole genome DNA methylation and found that while the ratio of 5-methylcytosine (5-mC) to total cytosine was similar between groups (Extended data Fig. 4a), the ratio of 5-hydroxymethylcytosine (5-hmC) modifications to total cytosine was significantly lower in csNOX4TG hearts (Extended data Fig. 4b). Integrating the set of differentially hydroxymethylated regions (DhMRs) located in gene bodies with our ATAC-seq dataset revealed that most of the regions were located in non-DAR areas, suggesting that these hydroxymethylation changes may play a role in regulatory mechanisms such as transcription elongation or alternative splicing rather than controlling gene accessibility (Extended data Fig. 4c). We also assessed the association between transcribed genes (RNA-seq dataset) and DhMRs and found that the majority of associated genes were hypohydroxymethylated (Extended Fig. 4d). The majority of the differentially expressed DhMR-associated genes (∼20% of all DhMR-associated genes) are not known targets of NRF2 or ATF4 (Extended data Fig. 4e-f). Functional enrichment analyses of these genes indicated several metabolic pathways linked to the overall changes in metabolism induced by NOX4 (Extended data Fig. 4g). Taken together, these results suggest that initial changes in glucose intermediary metabolism induced by NOX4 via the direct activation of a limited set of transcription factors may then trigger wider metabolic reprogramming through epigenetic changes, including a significant remodelling of histone and DNA methylation landscape (Fig. 4f).

The importance of metabolic reprogramming to support rapid cell proliferation in settings such as cancer or stem cell proliferation is well recognised. Notably, there is an increased activity of ancillary glycolytic pathways that support biomass generation, redox homeostasis, stress resistance and signalling.^13,16,17^ These include the PPP, which generates NADPH for reductive biosynthesis and redox homeostasis and ribose for nucleotide synthesis; the SBP, which generates serine and glycine for protein, nucleotide and glutathione synthesis; the dihydroxyacetone phosphate pathway, which contributes to triglyceride synthesis; and the HBP, which mediates protein glycosylation and signalling. Post-mitotic terminally differentiated cardiomyocytes in the adult heart have minimal proliferative capacity but do enlarge through hypertrophic remodelling under conditions of increased haemodynamic workload.^3^ The nature and extent of reprogramming of intermediary metabolism that the adult heart is able to undergo and how this may be regulated is unclear.^4,18^ A major finding of the present study is that the redox-signalling protein NOX4 (which is induced in response to haemodynamic overload) drives complex metabolic reprogramming in the adult terminally differentiated heart with features highly analogous to those reported in proliferating tissues. As such, we identify a significant redirection of glucose carbon flux towards the PPP, the SBP and the HBP, as well as evidence to support an increase in intermediates in other anabolic pathways such as glycerophospholipid synthesis and in the SGOC. Given the high energetic requirement of the adult heart, a relative redirection of glucose carbon flux away from the TCA cycle could significantly compromise contractile function if there are no other changes in energy production. However, in the case of NOX4-mediated metabolic reprogramming, these changes in glucose intermediary metabolism are accompanied by a significant increase in fatty acid oxidation with the consequence that cardiac energetic status is well preserved.^12^ Moreover, our prior work showed that the NOX4-mediated increase in fatty acid oxidation is mediated, at least in part, via increased O’GlcNAcylation of the fatty acid transporter CD36,^12^ indicating that the alterations in glucose intermediary metabolism may be mechanistically integrated with a change in fuel source for energy production. Collectively, this NOX4-induced metabolic reprogramming may position the heart to more effectively respond to and function during pathological stresses such as chronically increased haemodynamic workload, in line with our previous findings that NOX4 promotes compensated hypertrophic remodelling and function in this setting.^6^

We identify that a major component of the NOX4-induced metabolic programme appears to be transcriptionally driven in that most changes in metabolic pathways are mirrored by corresponding alterations in gene expression. Notably, we observe strong signatures for NRF2- and ATF4-regulated genes; the activation of both these transcription factors has previously been shown to be directly enhanced by endogenous NOX4.^7-10^ Known targets of NRF2 whose gene expression is found to be increased in NOX4-overexpressing hearts in the current study include key enzymes in glycolysis (*Hk1*), the PPP (*G6pd, Pgd, Tkt, Taldo1*), the SBP (*Phgdh*), glutathione metabolism/redox homeostasis (*Gss, Gclm, Gclc, Gsta2, Gpx1, Gsr*) and glycogen synthesis (*Gbe1*).^19^ Known ATF4 targets found to have increased gene expression include enzymes in the PPP (*G6pd*), SBP (*Phgdh, Psat1*), SGOC (*Shmt2*), HBP (*Gfat1, Gnpnat1*), and amino acid metabolism and translation (*Slc1a4, Slc7a11, Asns, Aars1*).^20^ Moreover, NRF2 and ATF4 may act in concert to regulate many metabolic pathways since ATF4 can heterodimerise with NRF2^21^ as well as act as a transcriptional and metabolic activator.^22^

The activation of ATF4 downstream of eukaryotic initiation factor 2α (eIF2α) phosphorylation has been shown to be specifically enhanced by NOX4 through a spatially-localized redox inhibition of protein phosphatase-1 (which reverses eIF2α phosphorylation), occurring at the endoplasmic reticulum (ER)^9^ – the major intracellular site where NOX4 is localised in cardiac and other cells.^23^ Previous work also showed that endogenous NOX4 has an obligatory role in the activation of NRF2 in the heart during physiological exercise, pathological haemodynamic overload or neurohumoral stimulation.^8,10^ In other words, there is a specific activation of NRF2 by NOX4 and not simply a non-specific oxidative effect; for example, the closely related ROS-producing NOX2 oxidase does not induce NRF2 activation in this setting.^8^ While the intracellular location of NOX4-dependent NRF2 activation was not investigated in the above study, it is interesting to note that NRF2 activation is reported to occur at the ER,^24^ suggesting that there may be coordinated activation of the two transcription factors at this site by NOX4.

Beyond ATF4 and NRF2, our analyses suggested multiple other transcription factors that putatively contribute to the overall NOX4-regulated metabolic rewiring. However, we considered it unlikely that NOX4 would specifically and directly activate so many transcription factors and therefore looked for alternative indirect mechanisms. Indeed, we find evidence of broad NOX4-induced epigenetic changes in histone and DNA methylation. The extensive remodelling of H3K4me3 marks was associated in csNOXTG hearts with genes involved in many of the metabolic pathways identified as being perturbed in our metabolomics analyses. We also found a perturbation of DNA hydroxymethylation (5-hmC marks) but no change in 5-mC marks. DNA hydroxymethylation has been shown to have dynamic effects on cardiomyocyte gene expression during development and hypertrophy through marks both on gene bodies and distal enhancer elements,^25^ and our results show it is linked to metabolic genes that are part of the overall NOX4-induced programme. The data therefore suggest that NOX4-induced epigenetic changes act in concert with more direct ATF4- and NRF2-mediated effects to generate the overall NOX4-dependent metabolic programme (Fig. 4f). Although the precise mechanisms underlying the extensive epigenetic regulation require further investigation, it seems likely that the initial NOX4-induced direct (NRF2- and ATF4-mediated) changes in glucose intermediary metabolism may then lead to epigenetic changes through altered availability of metabolites required for epigenetic modification and/or the activity of key enzymes involved in epigenetic modifications.^15^ For example, both histone demethylases and TET (ten-eleven translocation) enzymes (which facilitate the conversion from 5-mC to 5-hmC and further DNA demethylation) are α-ketoglutarate-dependent,^15,26^ while the SGOC product S-adenosyl methionine provides methyl groups for methylation reactions.^16^ However, this does not exclude the possibility that additional non-metabolic mechanisms may also be involved.

To our knowledge, this is the first report of the specific induction of Warburg-like metabolic rewiring in the terminally differentiated heart which can potentially support the dual needs of growth and contraction. Combined with our previous findings on NOX4-regulated changes in fatty acid oxidation, the current work shows that it is possible to simultaneously activate anabolic programmes reliant on glucose-derived carbons and rebalance energetic pathways to maintain optimal cardiac contractile function. The elucidation of mechanisms underlying this metabolic reprogramming may be relevant to the design of therapies to promote more physiological (i.e. compensated) cardiac remodelling during chronic disease stress.

## Acknowledgements

This work was supported by the British Heart Foundation (grants CH/1999001/11735, RG/20/3/34823 and RE/18/2/34213).

## Methods

### Animal and isolated heart studies

Procedures were conducted in compliance with the UK Home Office Guidance on the Operation of the Animals (Scientific Procedures) Act, 1986 and institutional ethical approval. Cardiomyocyte-specific Nox4-overexpressing mice (csNOX4TG) were described previously.^6^ All studies were performed in adult mice aged 3-4 months. Stable isotope-resolved metabolomics in isolated Langendorff-perfused hearts was performed as described.^26^ Hearts were first perfused with Krebs-Henseleit (KH) buffer containing 5 mM glucose, 0.4 mM palmitate, 0.5 mM octanoate, 1.5 mM lactate, 0.2 mM pyruvate, 0.5 mM glutamine, 100 IU/ml insulin, and 0.5 nM isoproterenol; then switched to a modified KH-buffer in which glucose was replaced by 5 mM U-^13^C_6_-glucose. Hearts were freeze-clamped for metabolic quenching and stored at -80°C. Glycogen content in frozen tissues (∼20 mg) was measured using a commercial assay according to the manufacturer’s instructions (Sigma-Aldrich).

### Liquid chromatography/mass spectrometry-based targeted metabolomics

Metabolite extraction from frozen heart tissue was performed as described.^27^ Polar metabolites were reconstituted in acetonitrile/water (v/v, 3/2) and injected into a 1290 Infinity II ultrahigh performance liquid chromatography (UHPLC) system coupled to a 6546-quadrupole time-of-flight (Q-TOF) mass spectrometer (Agilent Technologies) equipped with a dual AJS electrospray ionization source (Agilent Technologies).^27^ Chromatographic separation was performed with a Poroshell 120 HILIC-Z column. The mobile phase consisted of solution A (10 mM ammonium acetate, pH 9.0) and solution B (10 mM ammonium acetate/acetonitrile 15:85 (v/v), pH 9.0), both solvents containing 5 μM InfinityLab deactivator additive (5191-4506, Agilent Technologies). The elution gradient was: isocratic step at 95% B for 2 min, 95 to 65% B for 12 min, maintained at 65% B for 3 min, returned to initial conditions over 1 min, and then the column was equilibrated at initial conditions for 8 min. MS analyses were obtained in negative ionisation mode and dynamic mass axis calibration was achieved by continuous infusion post-chromatography of a reference mass solution using an isocratic pump connected to an ESI ionization source. MS acquisition rate was 1.5 spectra/sec and m/z data ranging from 50-1,200 were stored. Feature annotation and metabolite identification were based on accurate mass and standard retention times with a tolerance of ±5 ppm and ±0.5 min, respectively, and performed with MassHunter Profinder (v10.0.2; Agilent Technologies) using our in-house curated metabolite library based on metabolite standards (SigmaAldrich). ^13^C label incorporation levels were normalized to the natural occurrence of ^13^C isotopes and are represented as corrected abundance percentages. Comparisons of percentage ^13^C labelling were performed using Mixed Model Analysis on GraphPad Prism 10. Pathway enrichment analysis was performed using MetaboAnalyst 5.0 Software.^27^

### RNA sequencing and analysis

RNA was isolated from 20 mg homogenized, frozen heart tissue using the ReliaPrep™ RNA miniprep system (Promega, Z6112), following the protocol for fibrous tissue. After confirming RNA integrity (Agilent Bioanalyzer), libraries were prepared from 1 µg RNA per sample using the NEBNext® Ultra™ II Directional RNA Library Prep Kit with NEBNext® Poly(A) mRNA Magnetic Isolation Module (NEB, E7420S and E7490S), according to the manufacturer’s guidelines. Equimolar pooling of libraries for multiplexing was based on Qubit and Bioanalyzer analyses. Sequencing was performed on an Illumina NextSeq 500 platform with a 75 bp read length at our institutional core facilities. Differential gene expression (DEG) was identified using Partek® Flow software (Partek, St. Louis, MO, USA).

DEGs were subjected to Ingenuity Pathway Analysis (IPA, www.ingenuity.com) for metabolic and signalling pathways and upstream regulator identification. Significance thresholds were set at a -log(p-value) of >1.3, activation z-scores of >2 and inhibition z-scores of ≤2.

Gene set enrichment analysis was performed using the Broad Institute software, targeting the “C2: curated gene sets” from the Molecular Signatures Database (MSigDB v2022.1). This analysis incorporated gene sets from major canonical pathway databases, including BioCarta, KEGG, PID, Reactome, and WikiPathways.

### RT-qPCR

Quantitative RT-PCR was performed using the StepOnePlus system (Applied Biosystems) and qPCRBIO SyGreen Mix Hi-ROX (PCR Biosystems Ltd, Cat. PB20.12-05). Gene expression levels were normalized to *Hprt* using the comparative Ct method. Primer sequences are detailed in Supplementary Table 2.

### ChIP - qPCR

ChIP was carried out following standard protocols. Immunoprecipitation was performed using antibodies against NRF2 (Cell Signalling, mAb #12721) and ATF4 (Cell Signalling, mAb #11815). Following reversal of cross-linking and DNA purification, qPCR was performed using primers specific for the promoter regions of target genes. ChIP enrichment was calculated as fold enrichment relative to WT samples, after normalization to both the respective input control and the gene desert region. Statistical comparisons were performed using unpaired t-tests. The sequences of primers are given in Table 2.

### ATAC-seq

Nuclei were isolated from heart tissue using a modification of a published protocol.^28^ Specifically, 10 μM MG132 and 1 mM DTT were incorporated into the cell lysis buffer. Following homogenization, tissue samples were filtered through a 40 µm strainer and centrifuged at 600 g for 5 min. The resulting pellets were resuspended in 1.5 mL of cold cell lysis buffer and subsequently collected by centrifugation at the same speed. 50,000 nuclei were isolated from 1 mL of the resuspension in cold PBS and immediately subjected to ATAC-Seq using an established protocol.^29^ Library amplification used NEBNext High-Fidelity 2× PCR Master Mix and specific barcoded primers. After amplification, libraries were size-selected and quantified using AMPure XP beads and Agilent Bioanalyzer, respectively.

Sequencing was performed on an Illumina system, followed by data analysis that included mapping to the mm10 mouse genome, duplicate removal, and blacklisted region exclusion. Peak calling and differential accessibility analysis were executed using MACS2 and DESeq2. Differential accessibility regions (DARs) between the csNOX4TG and WT groups were evaluated by setting an FDR<0.1 and FC> 1.2 or <-1.2. For motif analysis within differential accessibility regions (DARs), we employed the SeqPos motif tool from the Cistrome toolbox (http://cistrome.org/ap/), targeting both *Mus musculus* and *Homo sapiens* motifs. We applied a stringent p-value cutoff of 0.001 against the comprehensive cistrome motif database. This approach facilitates the identification of transcription factor binding landscapes, enhancing our understanding of regulatory mechanisms in DARs.

### DNA methylation analyses

Genomic DNA was extracted from around 20 mg heart tissue from 3 csNOX4TG and 3 matched WT mice and purified using standard methods. The quantification of 5hmC in the genomic DNA was conducted using LC-MS/MS, adhering to a previously described protocol.^30^ Statistical analyses were performed on GraphPad Prism 10 using Student’s t-test.

Genomic DNA also underwent enrichment for hydroxymethylated DNA regions using the hydroxymethylated DNA immunoprecipitation (hMeDIP) technique, with sequencing and initial data processing handled by Arraystar Inc. (Rockville, Maryland) pipelines.^31^ Enriched regions (peaks) from the hMeDIP-Seq data were annotated using the UCSC Mouse mm10 genome reference (https://genome.ucsc.edu/). Differentially hydroxymethylated regions (DhMRs) that showed statistically significant variations between the csNox4 TG and WT groups were identified using the diffReps tool, applying a threshold of fold change (FC) ≥ 2 or ≤ -2 and an FDR of <0.01 to minimize false positives. DhMRs meeting these criteria were classified as either Hyper-DhMRs (FC ≥2) or Hypo-DhMRs (FC ≤ -2), depending on the direction of change. For functional annotation, DhMRs located within gene bodies were identified based on the UCSC RefSeq database (mm10 version). Genes associated with these DhMRs were cross-referenced with RNA-seq analysis results and subjected to IPA to explore potential pathways affected by differential hydroxymethylation patterns.

### CUT&Tag analysis of H3K4me3

CUT&Tag profiling of H3K4me3 was performed by Active Motif (Carlsbad, CA, USA) as a commercial service using their standard protocol. Briefly, nuclei from 3 WT and 4 csNOX4TG samples were permeabilised and incubated with an H3K4me3 antibody (Cat 61379, Active Motif) followed by a protein A–Tn5 transposase fusion. Targeted tagmentation was used to generate sequencing libraries, which were PCR amplified, purified, and sequenced on an Illumina platform. Sequencing reads were processed using Active Motif’s standard pipeline, including adapter trimming, alignment to the reference genome, removal of duplicate reads, peak and differential peak calling, and differential binding analysis which was performed using MACS2 and DiffBind with DESeq2 normalization. Differential regions between groups were identified using an FDR < 0.05 and fold change >±1.2. Peak locations were annotated using GREAT (great.stanford.edu), with promoter regions defined as within 3 kb of the transcription start site (TSS) and distal regions defined as >3 kb from the TSS.

### Statistical analyses

Comparisons between groups were performed by mixed model analysis for groups of unequal sizes or by unpaired t-test, as indicated in the figure legends. Analyses were performed using GraphPad Prism 10.

## Figure legends

**Extended Data Fig. 1.1.**
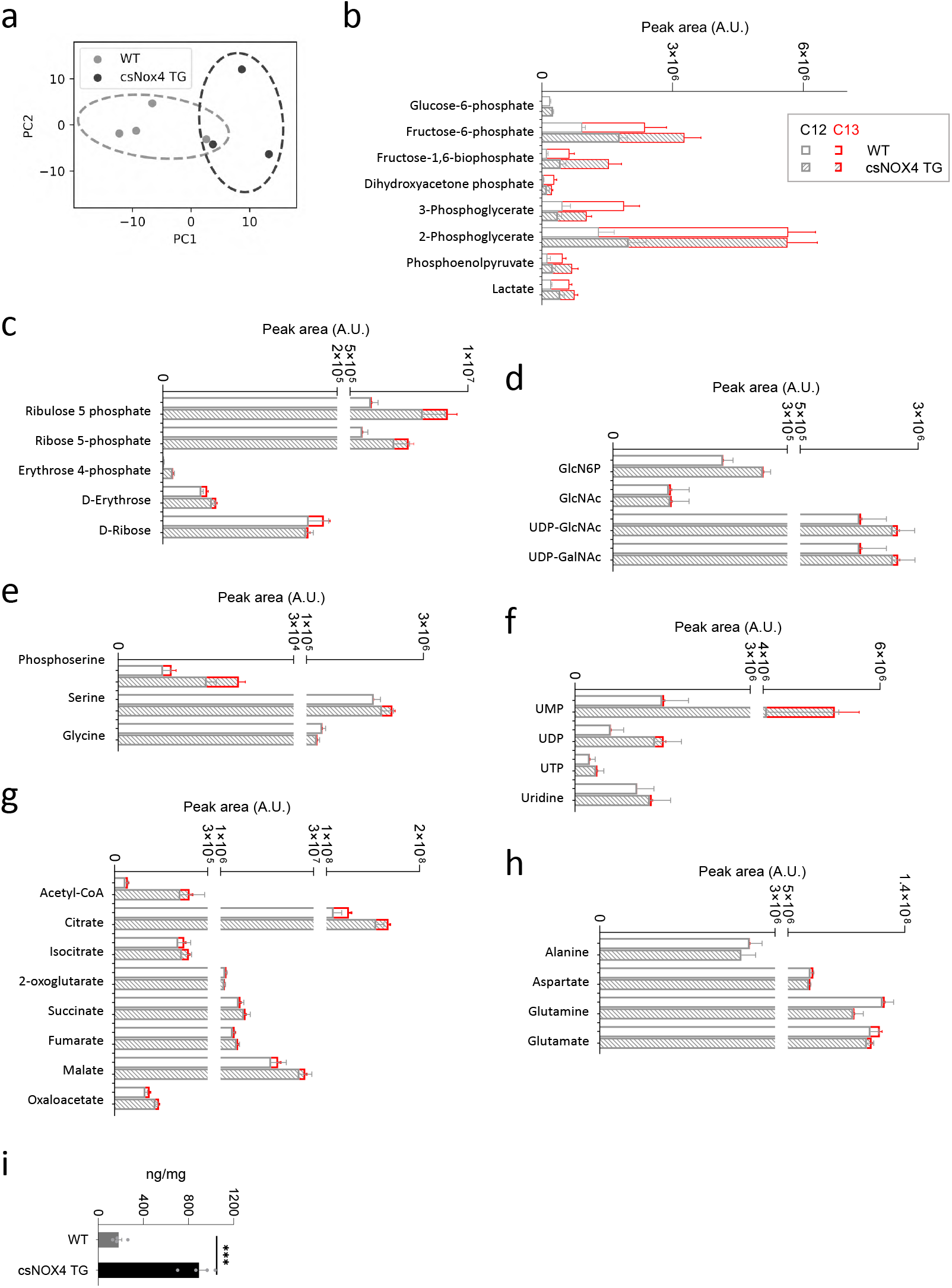
Effects of NOX4 on metabolites related to glucose intermediary metabolism in the heart. **a**, Principal component analysis (PCA) of cardiac metabolites in WT (n=3) and csNOX4TG (n=4) hearts. Analysis was performed using Python (scikit-learn, matplotlib). **b-h**, Comparative abundance (area under chromatographic peak) of labelled and non-labelled metabolites in WT and csNox4TG hearts for the experiment reported in Fig. 1d-j. Open bars show data for WT and filled bars for csNOX4TG. Data for ^13^C-labelled metabolites is depicted in red. **i**, Glycogen levels in WT and csNOX4TG hearts (n=4/group). ***p≤0.001.

**Extended data Fig. 2.**
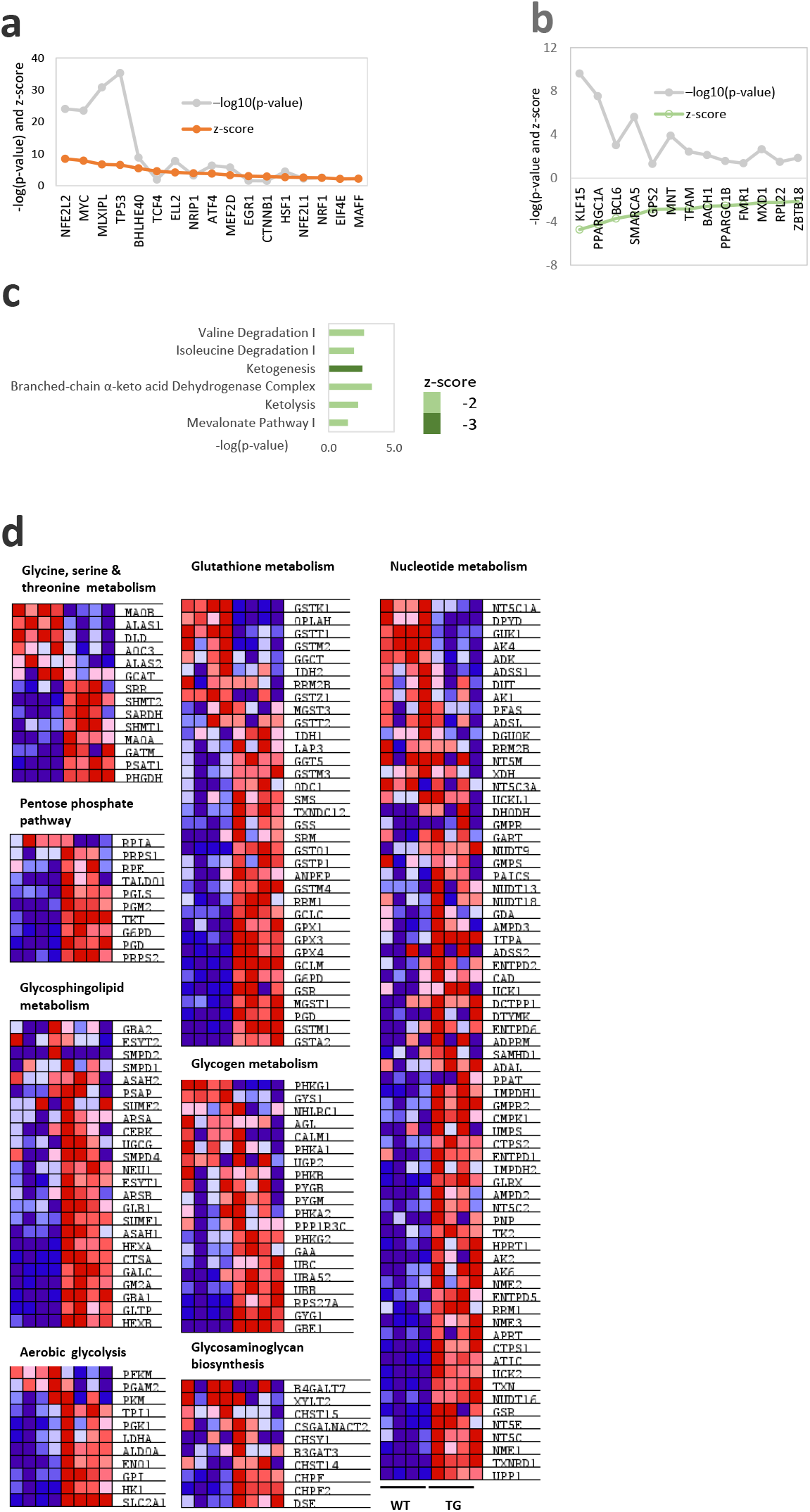
Predicted NOX4-dependent transcriptional regulators of gene expression. **a**, Top 15 predicted upstream transcription regulators that were activated. Significant enrichment cutoffs used were a z-score ≥ 2 and a -log(p-value) >1.3. **b**, Significantly repressed transcription regulators in csNOX4TG hearts. Cutoffs used were a z-score ≤ - 2, -log10 (p-value) >1.3. **c**, Significantly inhibited metabolic pathways enriched in csNOX4TG hearts, as analysed using IPA software. z-score of ≤ -2, -log (p-value) >1.3; *N*=4 biological replicates per group. **d**, Heatmap plots showing all the genes in the metabolic pathways analysed using GSEA in Fig. 2d-k. Dark red and dark blue represent greater levels of up- and down-regulation, respectively.

**Extended data Fig. 3.**
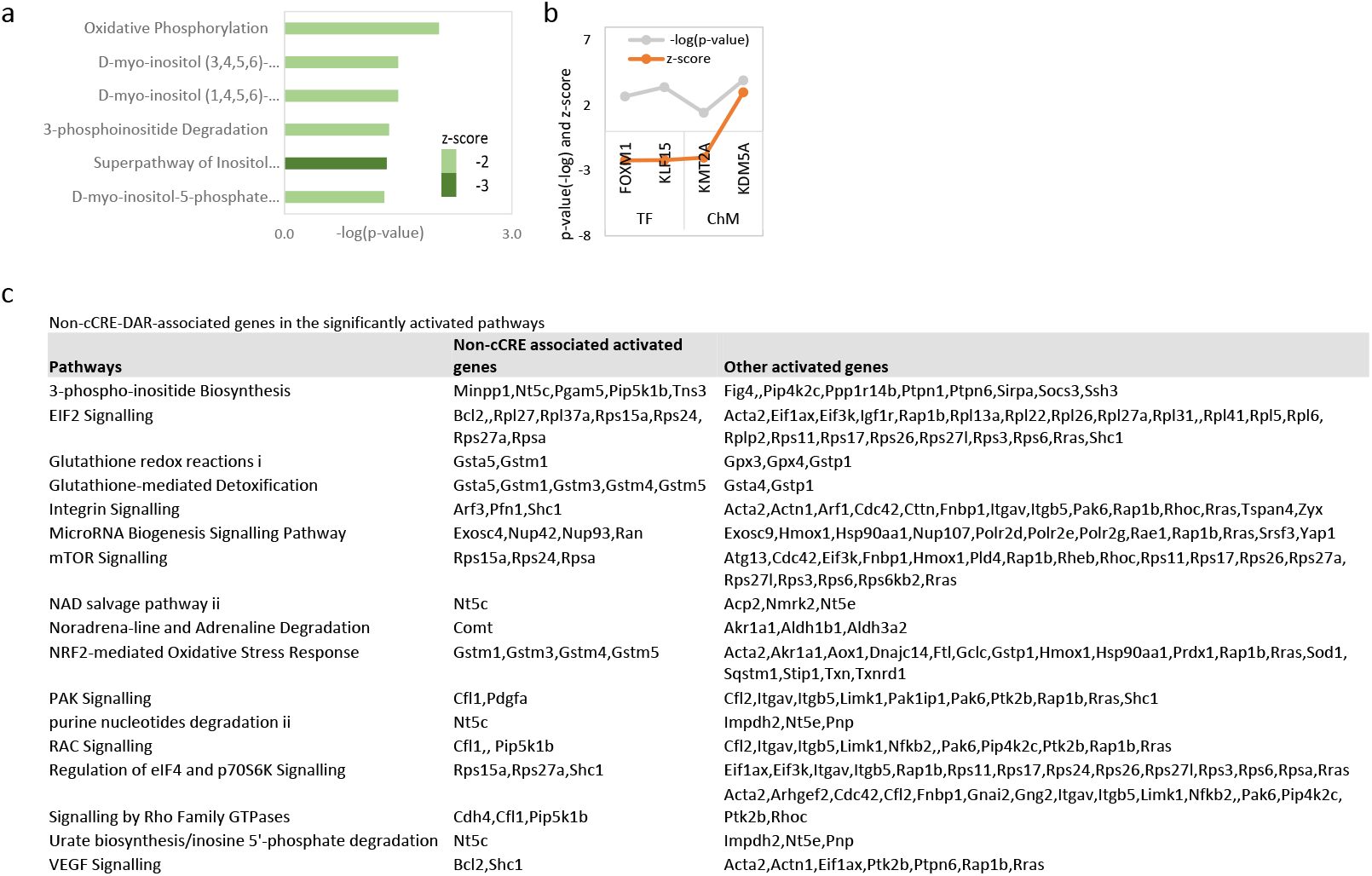
ATAC-seq analyses in csNOX4TG and WT hearts. **a**, IPA analysis of genes downregulated and associated with closed chromatin at DARs in TG hearts, for metabolic pathway enrichment. **b**, Predicted upstream transcriptional regulators for downregulated genes associated with closed chromatin at DARs in TG hearts. TF, transcription factor; ChM, chromatin modifier. **c**, List of genes associated with DARs and upregulated in significantly activated pathways in TG hearts.

**Extended data Fig. 4.**
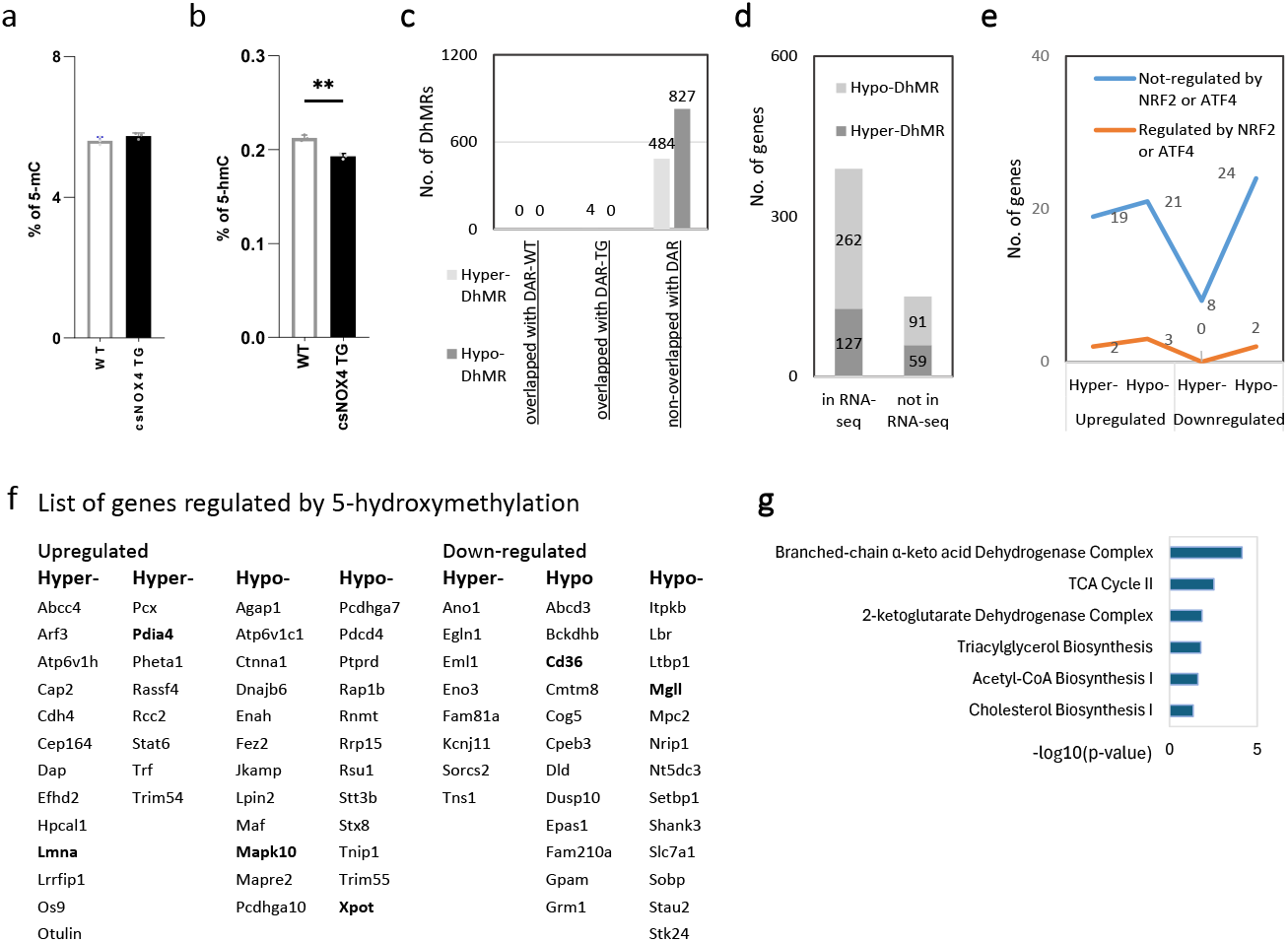
DNA methylation effects of NOX4. **a-b**, Ratio of 5-methylcytosine modifications (5-mC) to total cytosine and 5-hydroxymethylcytosine (5-hmC) to total cytosine in WT and csNOX4TG hearts (N=3/group). **p < 0.01. **c**, Numbers of hyper- or hypo-DhMRs on gene bodies and overlap with differentially accessible regions (DARs) on ATAC-seq. **d**, Distribution of differentially hydroxymethylated regions (DhMRs) in gene bodies of genes present or absent in the RNA-seq dataset. **e**, Differentially expressed genes (DEGs) associated with DhMRs. Orange line: genes regulated by NRF2 or ATF4; blue line: genes not regulated by these transcription factors. **f**, List of genes with altered 5-hydroxymethylation status and their differential gene expression in csNOX4TG hearts. Genes in bold are known to be regulated by NRF2 or ATF4 **g**, Functional pathway enrichment analysis of DhMR-associated DEGs.

**Supplementary table 1.** Fold change and statistical significance of metabolites grouped by metabolic pathway.

| Pathway |  |  |
| --- | --- | --- |
| Metabolite | FC (log<sub>2</sub>) | p-value (-log) |
| **Redox** |  |  |
| Glutathione (reduced) | 3.2 | 3.2 |
| Oxidized glutathione (GSSG) | 1.1 | 1.7 |
| S-Lactoylglutathione | 3.4 | 4.0 |
| **SGOC** |  |  |
| Adenosylhomocysteine | 1.3 | 1.4 |
| Cystathionine | 5.1 | 4.8 |
| Methylthioadenosine | 1.1 | 1.3 |
| N-Acetyl-methionine | 2.3 | 1.8 |
| N-Formylmethionine | 1.4 | 1.5 |
| **Glycolysis** |  |  |
| Glucose 6-phosphate | 2.9 | 1.4 |
| Fructose 6-phosphate | 2.0 | 1.4 |
| **PPP** |  |  |
| Ribose 5-phosphate | 2.0 | 1.6 |
| **HBP** |  |  |
| Glucosamine 6-phosphate | 1.0 | 1.4 |
| **SBP** |  |  |
| Phosphoserine | 4.7 | 2.1 |
| **Nucleotide** |  |  |
| AICAR | 3.1 | 2.7 |
| N-Carbamoyl-L-aspartic acid | 2.4 | 2.6 |
| Guanosine monophosphate | 1.7 | 1.5 |
| Guanosine diphosphate | 2.1 | 1.9 |
| UDP | 2.5 | 1.3 |
| **TCA Cycle** |  |  |
| cis-Aconitic acid | 1.2 | 2.1 |
| **Amino Acid** |  |  |
| Asparagine | 1.0 | 1.7 |
| Glutamic acid | 1.0 | 1.5 |
| L-Tyrosine | 1.8 | 1.8 |
| Phenylalanine | 1.7 | 1.5 |
| Proline | 1.6 | 1.6 |
| Tryptophan | 2.1 | 1.8 |
| Valine | 1.1 | 1.5 |
| **Modified Amino Acid** |  |  |
| Acetylcysteine | 4.1 | 3.2 |
| Acetylglutamic acid | 1.8 | 1.7 |
| Acetyllysine | 1.4 | 1.4 |
| **Others** |  |  |
| ADP-glucose | 1.0 | 1.3 |

**Supplementary table 2.**
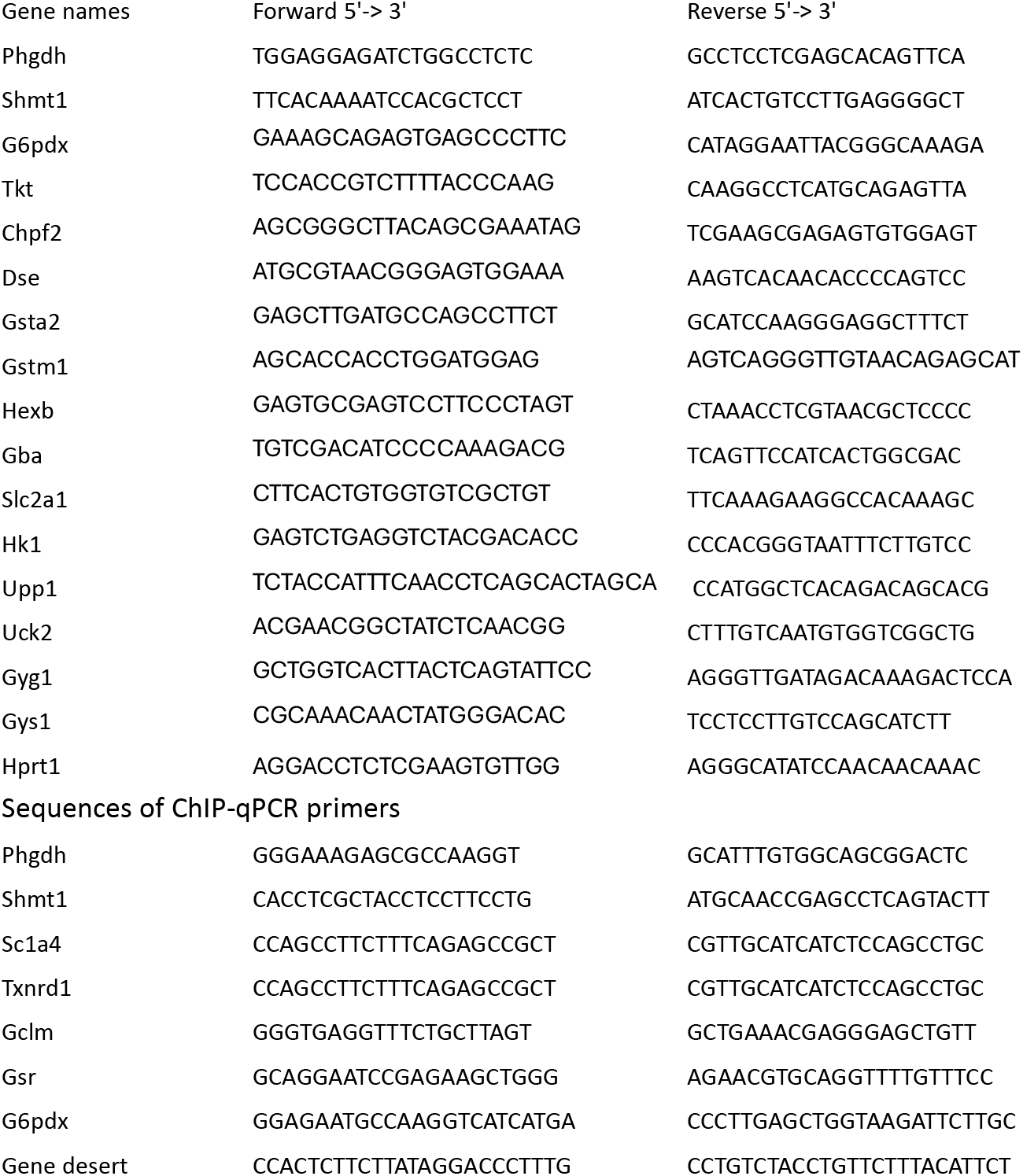
Sequences of RT-qPCR primers.

